# Mapping challenging mutations by whole-genome sequencing

**DOI:** 10.1101/036046

**Authors:** Harold E. Smith, Amy S. Fabritius, Aimee Jaramillo-Lambert, Andy Golden

**Affiliations:** National Institute of Diabetes and Digestive and Kidney Diseases, National Institutes of Health, Bethesda, Maryland 20892

**Keywords:** Forward genetics, variant detection, complex alleles, SNP mapping

## Abstract

Whole-genome sequencing provides a rapid and powerful method for identifying mutations on a global scale, and has spurred a renewed enthusiasm for classical genetic screens in model organisms. The most commonly characterized category of mutation consists of monogenic, recessive traits, due to their genetic tractability. Therefore, most of the mapping methods for mutation identification by whole-genome sequencing are directed toward alleles that fulfill those criteria (i.e., single-gene, homozygous variants). However, such approaches are not entirely suitable for the characterization of a variety of more challenging mutations, such as dominant and semi-dominant alleles or multigenic traits. Therefore, we have developed strategies for the identification of those classes of mutations, using polymorphism mapping in *Caenorhabditis elegans* as our model for validation. We also report an alternative approach for mutation identification from traditional recombinant crosses, and a solution to the technical challenge of sequencing sterile or terminally arrested strains where population size is limiting. The methods described herein extend the applicability of whole-genome sequencing to a broader spectrum of mutations, including classes that are difficult to map by traditional means.

## INTRODUCTION

The advent of next-generation sequencing technology has transformed our ability to identify the genetic variation that underlies phenotypic diversity. It is now feasible to perform whole-genome sequencing (WGS) in a matter of days and at relatively low cost. By comparing the data to an available reference genome, one can determine the complete constellation of sequence variants that are present in the sample of interest.

WGS has proven a boon to experimental geneticists who utilize classical, or forward, genetics (i.e., random mutagenesis and phenotypic screening) in model organisms. Although those techniques have been employed for more than a century (Morgan, 1911a; Morgan, 1911b), determination of the mutation responsible for the observed phenotype remains the rate-limiting step in most cases. Mutation identification by WGS compares favorably to traditional techniques, such as linkage mapping and positional cloning, in terms of speed, labor, and expense (Hobert, 2010; Bowerman, 2011; Hu, 2014). Consequently, the technology has been adopted in a wide variety of model species, including *Caenorhabditis elegans* (Sarin *et al*., 2008), *Drosophila melanogaster* (Blumenstiel *et al*., 2009), *Escherichia coli* (Barrick and Lenski, 2009), *Schizosaccharomyces pombe* (Irvine *et al*., 2009), *Arabidopsis thaliana* (Laitinen *et al*. 2010), *Saccharomyces cerevisiae* (Birkeland *et al*., 2010), *Dictyostelium discoideum* (Saxer *et al*., 2012), *Chlamydomonas reinhardtii* (Dutcher *et al*., 2012), and *Danio rerio* (Bowen *et al*., 2012).

A great advantage of forward genetic screening is the freedom from constraint on the types of alleles recovered; the only requirement is that the mutation produces the phenotype of interest. That property stands in contrast to techniques for reverse genetic screening, such as RNA interference (Fire *et al*., 1998), in which the molecular target is known but the ability to modify its activity is limited to a reduction of gene function. Although that category of mutation (reduction or loss of function) represents the class most commonly recovered in forward genetic screens, it is also possible to generate alleles that increase the level or activity of the gene product, reverse normal gene function, or produce novel activity. However, the expanded repertoire of variants accessible by random mutagenesis is accompanied by an increasing complexity in analysis. Although forward genetic screens are typically designed to obtain recessive alleles, they can also recover dominant, semi-dominant, and even multigenic mutations. The latter categories of alleles are challenging to map by traditional methods, such as linkage to morphological or molecular markers, due to the complexities of zygosity (for dominant and semi-dominant alleles) or the dependence of the phenotype upon independently segregating loci (for multigenic traits). As a practical consequence, preference is given to recessive, monogenic alleles, and the tools for mutation identification by WGS are tailored to that end.

We reasoned that the data produced by whole-genome sequencing, coupled with the appropriate mapping crosses, should greatly enhance our ability to identify causative variants among the less genetically tractable categories of mutations. To that end, we have utilized a polymorphism mapping method previously developed for mutation identification in *Caenorhabditis elegans* (Wicks *et al*., 2001; Swan *et al*., 2002; Doitsidou *et al*., 2010), and evaluated various strategies for mapping dominant, semi-dominant, and two-gene mutations. We have also performed analysis of WGS data from traditional recombinant mapping crosses, useful for mutation identification in legacy strains or in cases where polymorphism mapping may not be suitable. Finally, we have adopted a method to prepare sequencing libraries from small amounts of genomic DNA, and demonstrate its utility in strains where sample recovery may be limiting. Together, those approaches extend the application of WGS to the identification of various types of mutations.

## MATERIALS AND METHODS

All *C. elegans* mutant strains were derived from the wild-type N2 Bristol strain and contained one or more of the following alleles: *rol-6(su1006) II, lin-8(n111) II, lin-9(n112) III, spe-48(hc85) I* (formerly identified as *spe-8), spe-10(hc104) V, dpy-11(e224) V, unc-76(e911) V*. Hawaiian strain isolate CB4856 (Hodgkin & Doniach, 1997) was used for polymorphism mapping. CRISPR-mediated gene editing to create *spe-48(gd11)* was performed as described previously (Paix *et al*., 2015), using *dpy-10* as a co-CRISPR marker (Arribere *et al*., 2014). CRISPR reagents are listed in Supplement Table S1. Worms were propagated using standard growth conditions (Brenner, 1974). Fertility assays were performed as described previously (Kulkarni *et al*., 2012).

Sequencing libraries were constructed using the TruSeq DNA sample prep kit v2 for large input samples or TruSeq ChIP sample prep kit for low input samples (Illumina, San Diego, CA). Briefly, gDNA samples were sheared by sonication, end-repaired, A-tailed, adapter-ligated, and PCR-amplified. A detailed protocol for obtaining sheared gDNA from low input samples can be found in the Supplement (File S1). Libraries were sequenced on a HiSeq 2500 (Illumina, San Diego, CA) to generate 50bp reads, and yielded >20-fold genome coverage per sample. Genome version WS220 (www.wormbase.org) was used as the reference. The SNP data in Figure 1 were determined with a pipeline of BFAST (Homer *et al*., 2009) for alignment and SAMtools (Li *et al*., 2009) for variant calling. All other SNP data were obtained using BBMap (Bushnell, 2015) for alignment and FreeBayes (Garrison and Marth, 2012) for variant calling. Duplicate reads were removed after alignment and, unless otherwise indicated, at least three independent reads were required for a variant call. ANNOVAR (Wang *et al*., 2010) was used for annotation. For polymorphism mapping, SNP frequencies with LOESS regressions were plotted against chromosome position with R (R Core Team, 2015) using the same parameters reported for CloudMap (Minevich *et al*., 2012). Sequence data are available at the NCBI Sequence Read Archive (BioProject accession number PRJNA305991).

## RESULTS AND DISCUSSION

*SNP mapping strategy:* We employed a previously described one-step method for simultaneously mapping and identifying candidate mutations in *C. elegans* by WGS (Doitsidou *et al*., 2010). The parental strain bearing the mutation of interest is mated to a highly polymorphic Hawaiian strain isolate, which contains >100,000 annotated single-nucleotide polymorphisms (SNPs) (Wicks *et al*., 2001; Swan *et al*., 2002). Recombination between the parental and Hawaiian chromosomes occurs in the heterozygous F1 hermaphrodites, which are allowed to self-fertilize. Those F2 progeny that exhibit the mutant phenotype (mutant F2s, for short) are selected and pooled for sequencing. The positions and frequencies of the Hawaiian SNPs are plotted on the physical map; the SNPs are present in ~50% of the reads at unlinked loci, but are absent in the interval where the mutation resides (Figure 1, A-C). The same data are used to identify novel homozygous sequence variants within the mapping interval as candidate mutations. The strategy has been used successfully to identify a number of recessive mutations (e.g., Doitsidou *et al*., 2010; Labed *et al*., 2012; Liau *et al*., 2013; Connolly *et al*., 2014; Wang *et al*., 2014; Jaramillo-Lambert *et al*., 2015). With the exception of the strains that contain linked morphological markers (see below), we utilized the Hawaiian SNP mapping method described here for the characterization of different categories of mutations.

*Dominant and semi-dominant alleles:* For recessive alleles, homozygosity in the mutant F2s produces a gap in the Hawaiian SNPs that defines the mapping interval. Dominant alleles, however, yield a mixture of heterozygous and homozygous mutant F2s in a 2:1 ratio. Consequently, the frequency of Hawaiian SNPs at the mutant locus is only slightly reduced, from 50% to 33%, with a concomitantly small increase in the candidate allele frequency to 67%. Therefore, neither the mapping interval nor the causative mutation is clearly defined.

**Figure 1.**
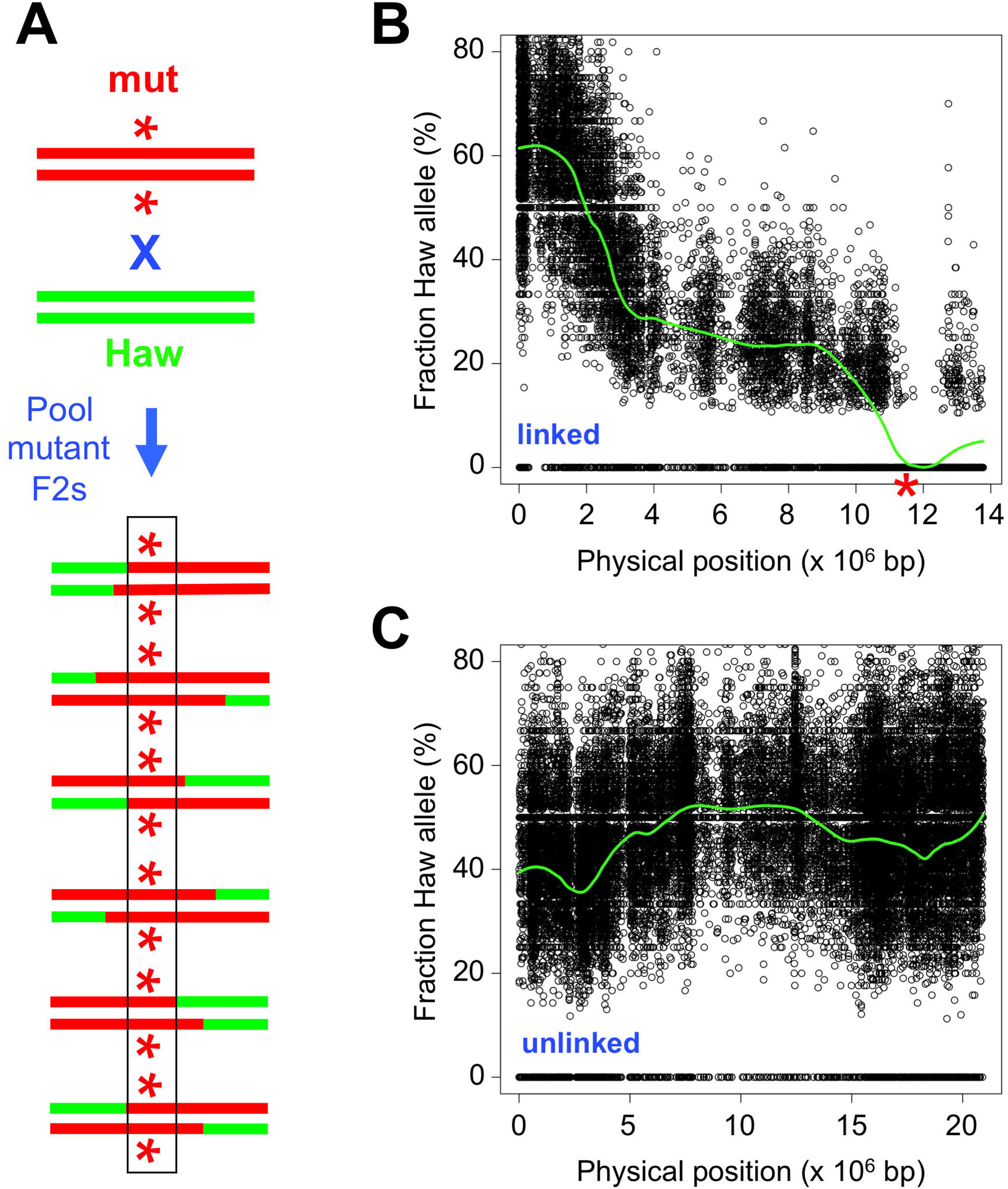

We considered two different strategies to address the problem (diagrammed in Figure 2A). For the first strategy, reverse mapping, wild-type F2s are picked and pooled for sequencing. The dominant allele is excluded from the population, so the Hawaiian SNP frequency rises to 100% in the mapping interval. An additional sequencing sample, from the mutation-bearing parental strain, is required to identify candidate mutations in that region. The second strategy involves the screening of F3 progeny from individual mutant F2s, to discriminate homozygous (100% mutant F3s) from heterozygous (75% mutant F3s) F2 animals. The homozygotes can then be pooled, sequenced, and analyzed in the same manner as recessive mutations.

We tested the two mapping strategies using the well-characterized *rol-6(su1006)* mutation. The *rol-6* gene encodes a cuticular collagen, and the *su1006* allele produces an easily identifiable dominant roller (Rol) phenotype (Kramer *et al*. 1990). For the reverse mapping strategy, we picked 100 wild-type F2 progeny from the mapping cross. The mapping plot produced an elevated frequency of SNPs on a single chromosome (chrII) with a peak that encompasses *rol-6* (Figure 2B). However the mapping interval was not as clearly demarcated as for recessive alleles, because the latter approach benefits from a threshold effect for SNP detection. Therefore, we also plotted the normalized frequency of homozygous SNPs in 0.5 megabase (Mb) intervals (Supplement Figure S1). We observed that the peak SNP frequency corresponds with the location of *rol-6*. For the F3 screen, we picked 120 individual F2 rollers and identified 36 homozygotes that produced 100% Rol progeny. Polymorphism mapping revealed a reduction in SNP frequency on chromosome II, with a gap that encompasses *rol-6* (Figure 2C). Our results demonstrate the validity of the two methods for mapping dominant mutations.

**Figure 2.**
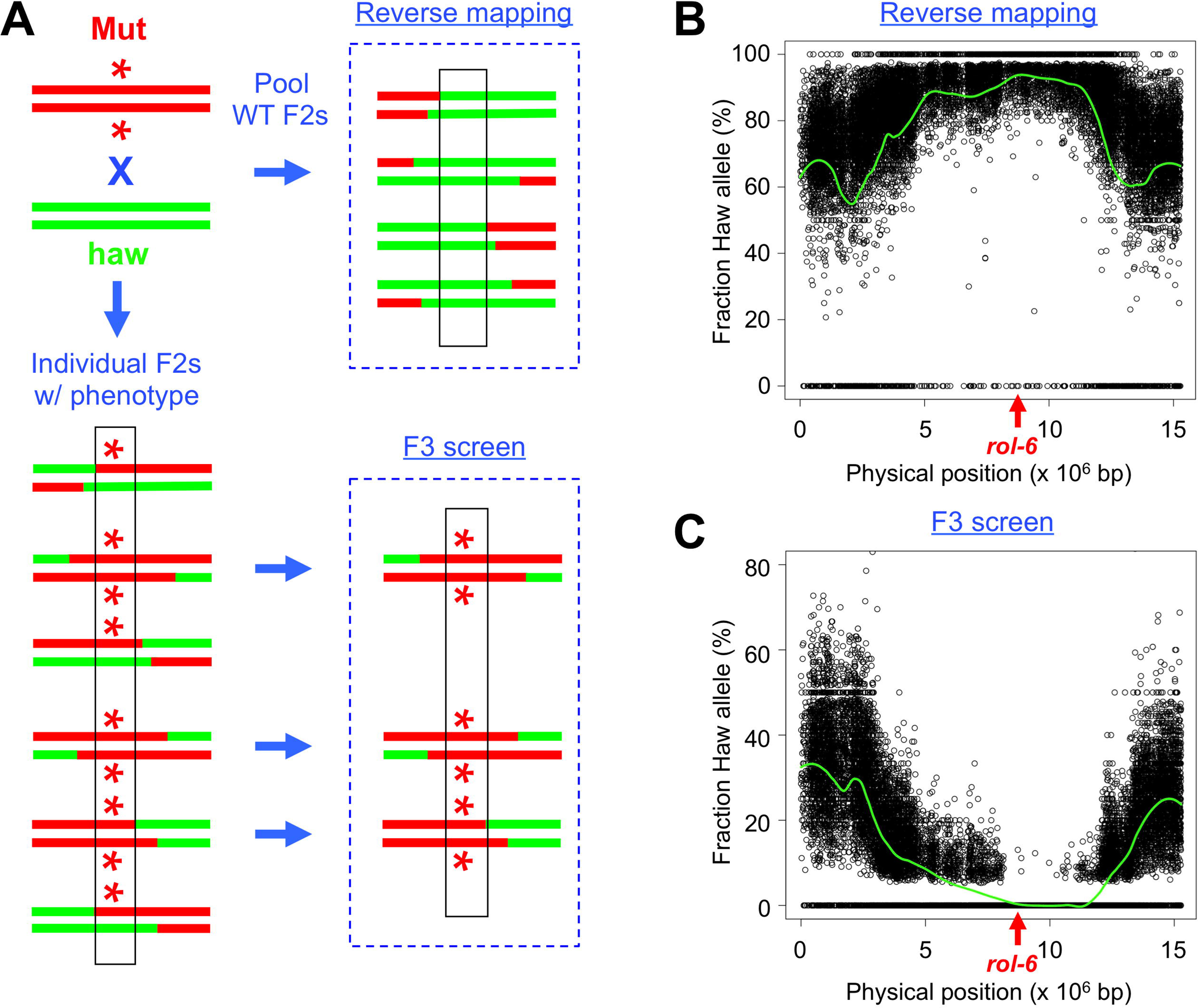

In principle, semi-dominant alleles should be more amenable to analysis than dominant alleles, since heterozygous animals exhibit an intermediate phenotype that can be distinguished from homozygotes. In practice, many phenotypes (such as body length or lifespan) are phenomena with a continuous range of values, and it may not be possible to assign the genotype of individuals unambiguously based on the observed characteristic. Therefore, we recommend the same F3 progeny screening strategy used for dominant alleles, which should prove similarly sufficient to distinguish the zygosity of individual F2 animals. The reverse mapping strategy might also be employed, but only in cases where the wild-type and heterozygous phenotypes are fully resolved.

*Two-gene synthetic interactions:* The category is defined by the requirement for mutations at two loci to produce the phenotype of interest. Although more typically isolated as genetic enhancers, it is also possible to obtain such double mutants by unbiased forward screening. One of the earliest screens in *C. elegans* for cell lineage mutants recovered the recessive *lin-8(n111); lin-9(n112)* pair of alleles that define the synthetic multivulva (synMuv) class A and B categories (Ferguson & Horvitz, 1989).

Each of those mutations is phenotypically wild-type in isolation, but the double mutant exhibits multiple ectopic pseudovulvae (Muv phenotype).

We chose the *lin-8; lin-9* pair as a test case for the application of WGS to the identification of two-gene synthetic interactors. We selected 100 F2 Muv progeny from the mapping cross and pooled them for sequencing. Two chromosomes contained a biased distribution of SNPs, with mapping intervals that correspond to the positions of *lin-8* and *lin-9* (Figure 3, A-C). The gap in SNPs flanking *lin-8* is masked by a high local density of polymorphisms, but evident at higher resolution (Figure 3B). We conclude that WGS and polymorphism mapping can be applied successfully to map two-gene traits.

**Figure 3.**
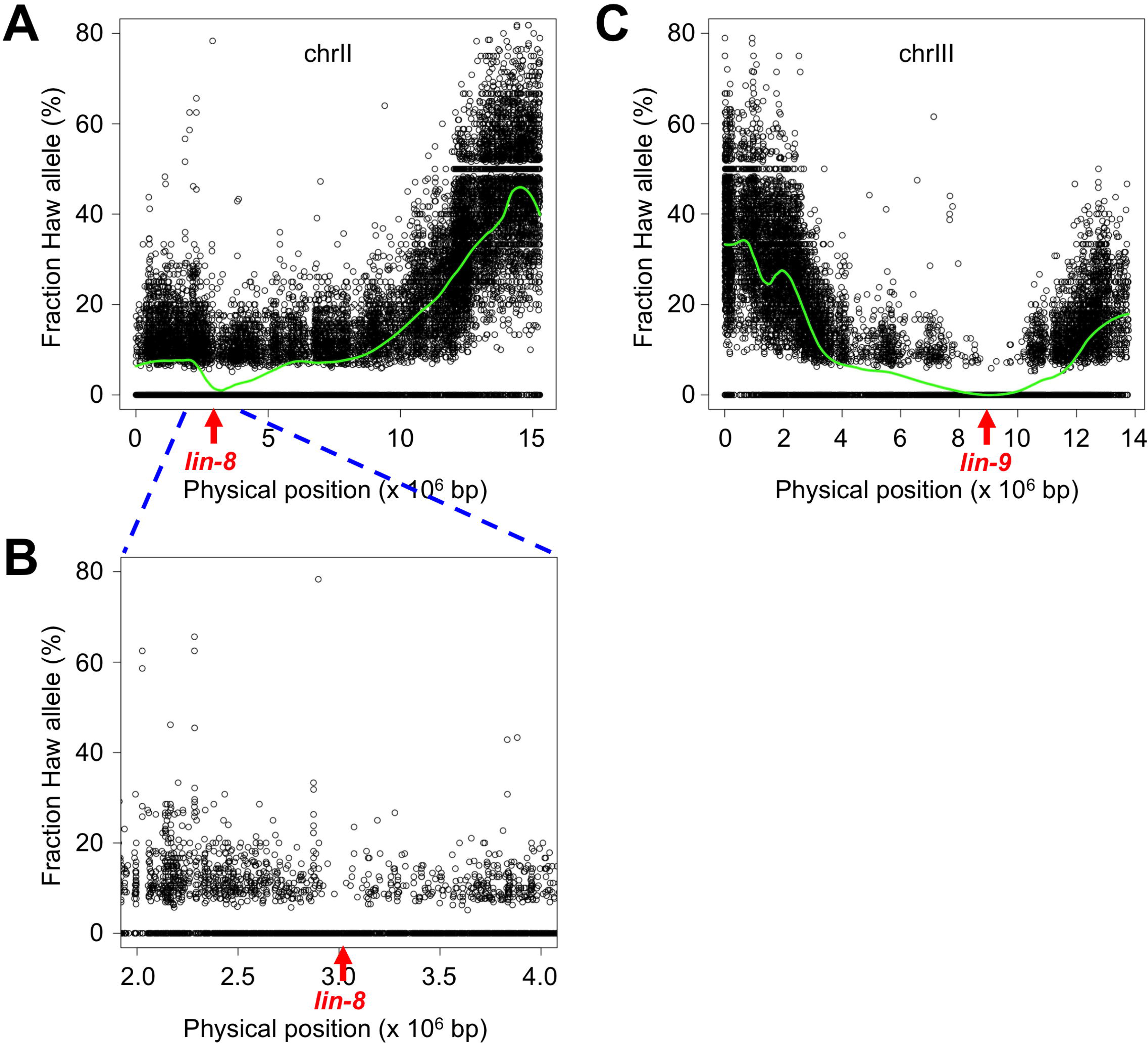

*Non-Hawaiian mapping:* The examples described above require mating to the polymorphic Hawaiian strain to identify the mapping interval. However, there are numerous cases where that cross may not be suitable. First, there are known genetic incompatibilities between the Hawaiian strain and the standard laboratory N2 Bristol strain (Seidel *et al*., 2008). The incompatibility loci produce biased segregation ratios in the F2 population, such that mutations in the vicinity of the loci are difficult to map (Minevich *et al*., 2012). Second, the genetic diversity of the polymorphic strain might include modifiers of the phenotype of interest, thereby confounding the isolation of phenotypically identifiable animals needed to generate the mapping information. Finally, the investigator may wish to take advantage of available recombinant lines obtained from crosses to morphologically marked strains. Those include legacy strains that were mapped using two or three-factor crosses, as well as lines constructed to link an easily visible marker to the mutation of interest.

A map-free method of mutation identification has been developed (Zuryn *et al*., 2010), in which homozygosity of linked variants after extensive backcrossing defines the mutation interval. An extension of that approach, using bulk segregant analysis, was subsequently reported (Abe *et al*., 2012; Minevich *et al*., 2012). We adopted a conceptually similar strategy for the analysis of recombinant lines from traditional mapping crosses (Figure 4A). Consider the simple case of a mutation linked to a morphological marker, obtained from a single recombinant founder. The line contains two sources of genetic variation that are absent from the wild-type strain: 1) *de novo* variants produced during mutagenesis, and 2) variants present in the morphologically marked strain that are introduced by recombination. The mapping interval can be determined from the novel variants by a single backcross to the wild-type strain. Screening for the marked mutant F2s imposes homozygosity on variants in the interval between the marker and mutation. Those animals are pooled, sequenced, and compared to the wild-type sample to identify novel variants. A cluster of novel homozygous variants on the physical map delimits the mapping interval. Note that the strategy is not predicated on knowledge of the physical location or molecular identity of the linked morphological marker allele, nor does it require accurate determination of the recombination frequency between the marker and the mutation of interest.

We evaluated the strategy using the *spe-10(hc104)* mutation marked with *dpy-11(e224)*. The former confers temperature-sensitive, sperm-specific sterility (Spe phenotype), and the latter produces short, fat animals (Dpy phenotype). We crossed the *dpy-11 spe-10* strain to wild-type males, and picked Spe Dpy F2s grown at the restrictive temperature for sequencing. After variant calling, we removed those that were also present in the wild-type sample and identified novel homozygous variants (minimum 15 reads, >80% variant call). The 80% threshold accommodates errors in the sequence data as well as mistakes in selecting the desired F2s, at the expense of including some non-homozygous variants. The distribution of homozygous variants was clearly enriched on chromosome V (73 of 87 total, or 84%). We plotted those variants and highlighted the *bona fide* (100%) homozygous calls (Figure 4B, in red), and observed a cluster between 5.0-14.5 Mb (a span that encompasses *dpy-11* at 6.5 Mb and *spe-10* at 10.4 Mb). We conclude that, in the case of strains bearing marked mutations, a single backcross is sufficient for mapping.

The same mapping method can be employed in cases where reciprocal recombinants are available from three-factor crosses with flanking markers. Individual homozygous mutant lines containing either the left or right marker are sequenced separately, and the frequencies of novel variants in each sample are plotted on the physical map. A contiguous cluster of homozygous variants present in both samples defines the mutation interval. The endpoints are delimited by variants that are homozygous in only one sample.

We assessed the utility of reciprocal mapping with a *spe-10(hc104) unc-76(e911)* strain (the latter mutation produces uncoordinated, or Unc, movement). After mating with wild-type males, we picked Spe Unc F2s grown at 25°C for sequencing. First, we mapped homozygous variants as above. The plot revealed a biased distribution on chromosome V (52 of 69 total variants, 75%), with a cluster between 8.0-16.5 Mb that contains *spe-10* and *unc-76* (located at 15.1 Mb; Figure 4C). For reciprocal mapping, we used the homozygous variants identified in either the *dpy-11 spe-10* sample or the *spe-10 unc-76* sample, and calculated the average variant fraction by combining the data from both samples. The map plot produced a smaller interval than either single sample, with well-defined endpoints at 8.6 and 14.5 Mbp clearly indicated by the LOESS curve (Figure 4D). The utility of the strategy should be weighed against the effort and expense of sequencing an additional strain, but it may be worthwhile when the markers are far from the mutation of interest and/or validation of candidate alleles is challenging.

**Figure 4.**
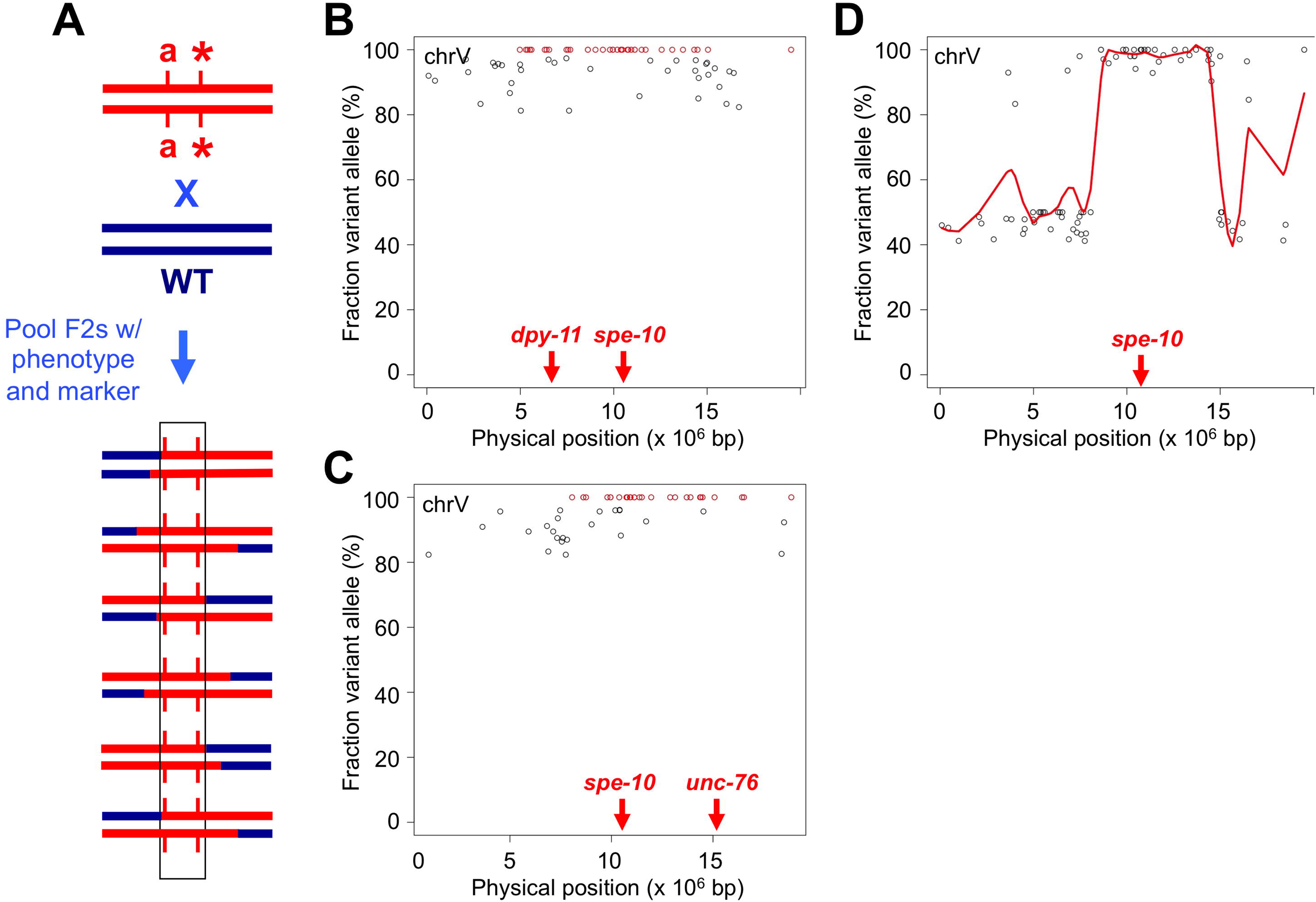

*Sequencing from small samples:* Many categories of mutations produce terminal phenotypes (such as lethality or sterility) that effectively limit the sample size. Standard WGS library preparation calls for microgram quantities of input gDNA, which can only be obtained from large populations. However, protocols for chromatin immunoprecipitation sequencing (ChIP-Seq) libraries require only low nanogram amounts of input, so we evaluated the suitability of those methods for WGS from small numbers of worms.

One concern is that the sequence coverage obtained by using ChIP-Seq library preparation might prove insufficient for our application. Typically, ChIP samples contain only a fraction of the genome, whereas accurate mutation identification requires data that encompass the entire genome. The potential for coverage bias from small amounts of input DNA might preclude detection of some variants. To address that concern, we compared the data produced from standard library preparation using 1 μg of gDNA (referred to as the large sample) to those generated from a ChIP-Seq library prep protocol using only 2 ng (hereafter, small sample) of the same gDNA. We selected the highly polymorphic Hawaiian strain as our test case, to ensure a large number of validated SNPs for identification. We used equal numbers of sequence reads for the two data sets and compared the alignment metrics.

First, we determined the fraction of the genome that was covered by at least three reads (the minimum read depth required for a variant call in our analysis) in the large and small samples. We observed little difference (<0.25%) in genome coverage between the two libraries (Figure 5A), indicating that the small sample does not introduce substantial gaps in coverage. Note that our coverage calculations are conservative; repetitive sequences constitute ~5% of the genome and, by filtering reads that mapped to multiple loci, those sequences were excluded. Next, we considered more subtle evidence for biased coverage. The small amount of input DNA might reduce the complexity of that library relative to the large sample, producing an increase in the fraction of duplicate reads as well as a larger variance in the per-base depth of coverage. We did observe differences in the degree of read duplication (large, 15.8%; small, 39.1%) as well as the depth of coverage distribution between the two libraries (Figure 5B), consistent with reduced library complexity in the small sample. However, the impact of those differences on SNP detection was minimal. After removing the duplicate reads from each sample, we performed variant calling and compared the lists of annotated Hawaiian SNPs that were identified in each data set. The numbers and identities of the SNP calls were essentially the same between the large and small samples, with greater than 99% overlap (Figure 5C). We conclude that libraries constructed from small amounts of input are sufficient for accurate mutation identification.

**Figure 5.**
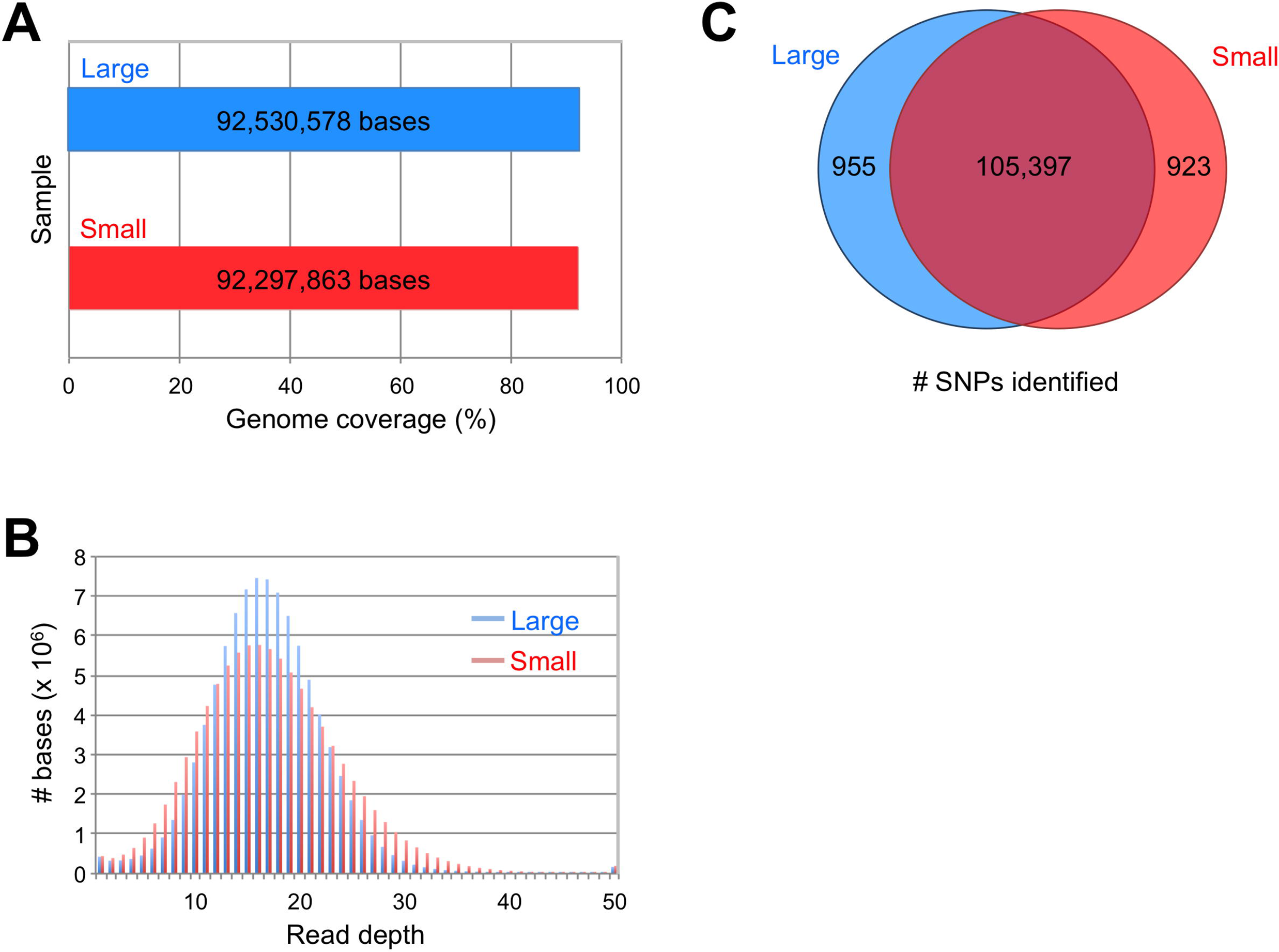

We proceeded to obtain gDNA from a small population of worms with a terminal mutant phenotype. Our test strain contained the recessive *hc85* allele, which produces a sperm-specific defect that renders them sterile (L’Hernault *et al*., 1988). The mutation was originally assigned to the *spe-8* complementation group. However, more recent data dispute that assignment; *hc85* complements the sperm-specific sterility of another *spe-8* allele, and targeted sequencing of the *spe-8* gene from the strain bearing *hc85* failed to identify a molecular lesion (Muhlrad *et al*., 2014). To address that discrepancy, we used the polymorphic mapping cross and recovered gDNA from a pool of 50 hand-picked sterile F2 animals. SNP mapping defined the interval between positions 1.7-2.7 Mb on chromosome I (Figure 6A). That interval is distinct from the position of *spe-8*, which is located on the same chromosome at position 0.1 Mb.

We confirmed the validity of the SNP mapping data by determining the molecular identity of the *hc85* allele. Four genes within the mapping interval contained novel, homozygous, nonsynonymous mutations (Figure 6B). Previous analyses determined that virtually all Spe genes exhibit sperm-specific expression (e.g., see Kulkarni *et al*., 2012), so the four candidates were prioritized by that criterion. Recently published RNA-Seq data (Ma *et al*., 2014) indicated that only one of those four genes, *Y51F10.10*, is detectably expressed in sperm. Therefore, we engineered the observed *Y51F10.10* missense mutation, a threonine-to-isoleucine substitution at amino acid 272, into the wild-type strain via CRISPR-mediated gene editing (Paix *et al*., 2015). The engineered allele, designated gd11, recapitulates the recessive Spe phenotype of *hc85*. Self-fertility is normal for heterozygous *gd11/+* hermaphrodites but low for homozygous *gd11* hermaphrodites, and fertility is restored by mating to wild-type males (Figure 6C). Complementation testing of the Spe phenotype revealed the *gd11* and *hc85* mutations to be allelic, as self-fertility of *gd11/hc85* hermaphrodites is low (Figure 6C). Because the *Y51F10.10* gene is distinct from *spe-8*, we have assigned the new gene name *spe-48* to *hc85* and *gd11*.

**Figure 6.**
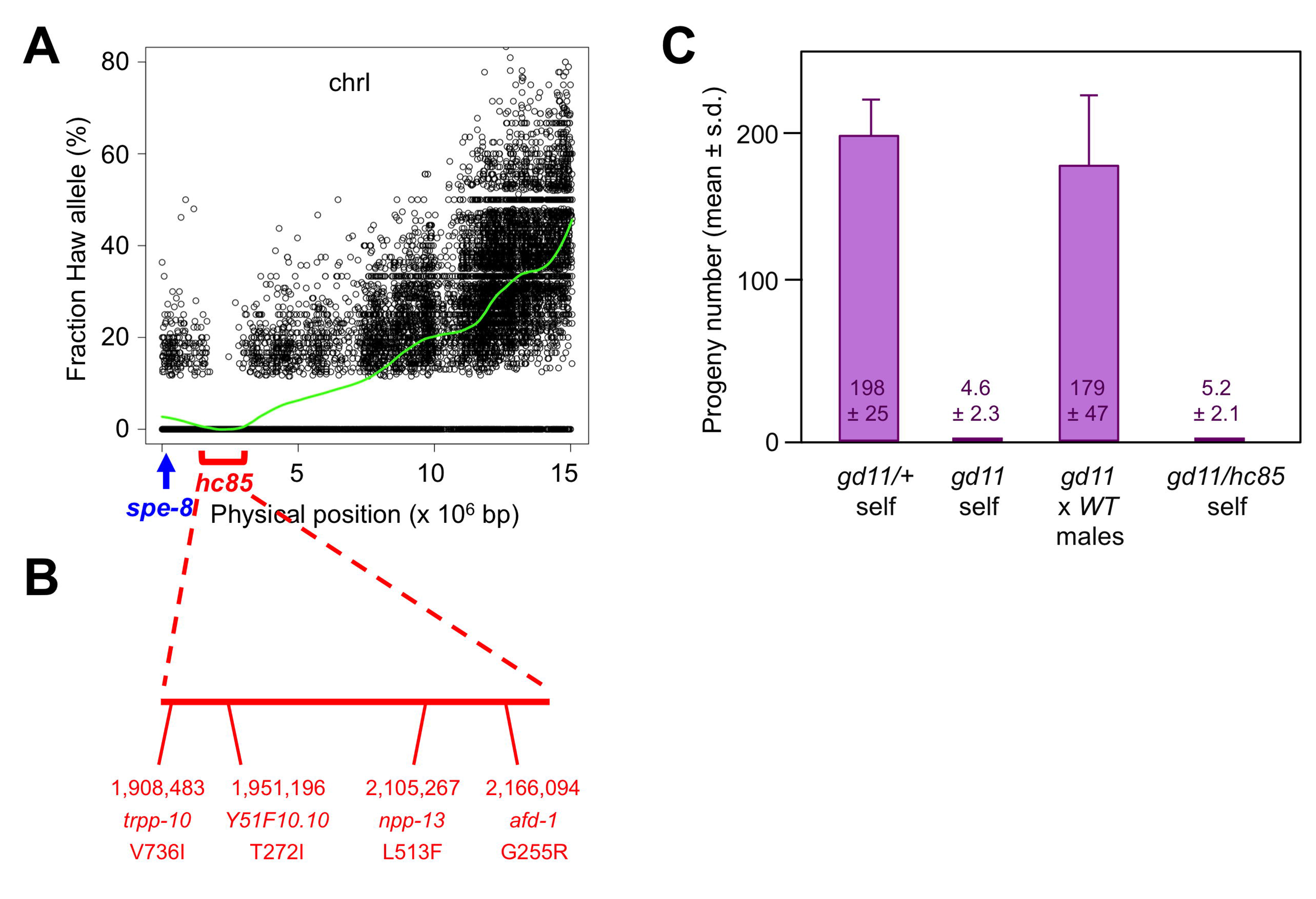

## CONCLUSION

Whole-genome sequencing has transformed our ability to identify mutations obtained from forward screening. We describe a variety of methods for applying the technology to types of mutations – dominant and semi-dominant alleles, synthetic interactors, and terminal phenotypes – that pose particular challenges to identification. We demonstrate the validity of those methods by the identification of both previously known as well as new mutations, and confirm the latter by independent criteria. Although our analyses were limited to *C. elegans*, the strategies can be generalized to any species in which polymorphic isolates are available for crosses and should prove useful for researchers who wish to apply the power of whole-genome sequencing to the investigation of alleles that are difficult to identify by other methods.

## ACKNOWLEDGMENTS

We thank members of Michael Krause’s lab and the Baltimore Worm Club for fruitful discussions, and Sevinc Ercan for sharing the small sample gDNA isolation protocol. Some strains were provided by the Caenorhabditis Genetics Center, which is funded by the NIH Office of Research Infrastructure Programs (P40 OD10440). This study was supported by the Intramural Research Program of the National Institutes of Health, National Institute of Diabetes and Digestive and Kidney Diseases, and is subject to the NIH Public Access Policy.

